# Structural remodeling of SARS-CoV-2 spike protein glycans reveals the regulatory roles in receptor binding affinity

**DOI:** 10.1101/2021.08.26.457782

**Authors:** Yen-Pang Hsu, Debopreeti Mukherjee, Vladimir Shchurik, Alexey Makarov, Benjamin F. Mann

**Affiliations:** Analytical Research and Development, Merck & Co., Inc., Rahway, NJ 07065, USA

## Abstract

Glycans of the SARS-CoV-2 spike protein are speculated to play functional roles in the infection processes as they extensively cover the protein surface and are highly conserved across the variants. To date, the spike protein has become the principal target for vaccine and therapeutic development while the exact effects of its glycosylation remain elusive. Experimental reports have described the heterogeneity of the spike protein glycosylation profile. Subsequent molecular simulation studies provided a knowledge basis of the glycan functions. However, there are no studies to date on the role of discrete glycoforms on the spike protein pathobiology. Building an understanding of its role in SARS-CoV-2 is important as we continue to develop effective medicines and vaccines to combat the disease. Herein, we used designed combinations of glycoengineering enzymes to simplify and control the glycosylation profile of the spike protein receptor-binding domain (RBD). Measurements of the receptor binding affinity revealed the regulatory effects of the RBD glycans. Remarkably, opposite effects were observed from differently remodeled glycans, which presents a potential strategy for modulating the spike protein behaviors through glycoengineering. Moreover, we found that the reported anti-SARS-CoV-(2) antibody, S309, neutralizes the impact of different RBD glycoforms on the receptor binding affinity. Overall, this work reports the regulatory roles that glycosylation plays in the interaction between the viral spike protein and host receptor, providing new insights into the nature of SARS-CoV-2. Beyond this study, enzymatic remodeling of glycosylation offers the opportunity to understand the fundamental role of specific glycoforms on glycoconjugates across molecular biology.

**Covert art Legends:** The glycosylation of the SARS-CoV-2 spike protein receptor-binding domain has regulatory effects on the receptor binding affinity. Sialylation or not determines the “stabilizing” or “destabilizing” effect of the glycans. (Protein structure model is adapted from Protein Data Bank: 6moj. The original model does not contain the glycan structure.)

**Significance:** Glycans extensively cover the surface of SARS-CoV-2 spike (S) protein but the relationships between the glycan structures and the protein pathological behaviors remain elusive. Herein, we simplified and harmonized the glycan structures in the S protein receptor-binding domain and reported their regulatory roles in human receptor interaction. Opposite regulatory effects were observed and were determined by discrete glycan structures, which can be neutralized by the reported S309 antibody binding to the S protein. This report provides new insight into the mechanism of SARS-CoV-2 S protein infection as well as S309 neutralization.

## Introduction

Host cell infection of severe acute respiratory syndrome coronavirus 2 (SARS-CoV-2) is initiated by the interaction between the viral spike (S) proteins and the host cellular receptors, angiotensin-converting enzyme 2 (ACE2).(1, 2) The S protein recognizes ACE2, mediates protease priming and then promotes virus membrane fusion.(3) Because of its critical roles in the infection processes, the S protein has become the principal target for vaccine and therapeutic development.(4, 5) Similar to other β-coronavirus members, the SARS-CoV-2 S protein is heavily glycosylated.(6, 7) It contains 22 N-linked glycosylation sites with 18 of them conserved from the SARS-CoV spike **(Figure 1A)**.(7, 8) The glycan moieties contribute about 17% molecular weight to the native-state trimetric SARS-CoV-2 S protein, shielding approximately 40% of the protein surface.(9)

**Figure 1.**
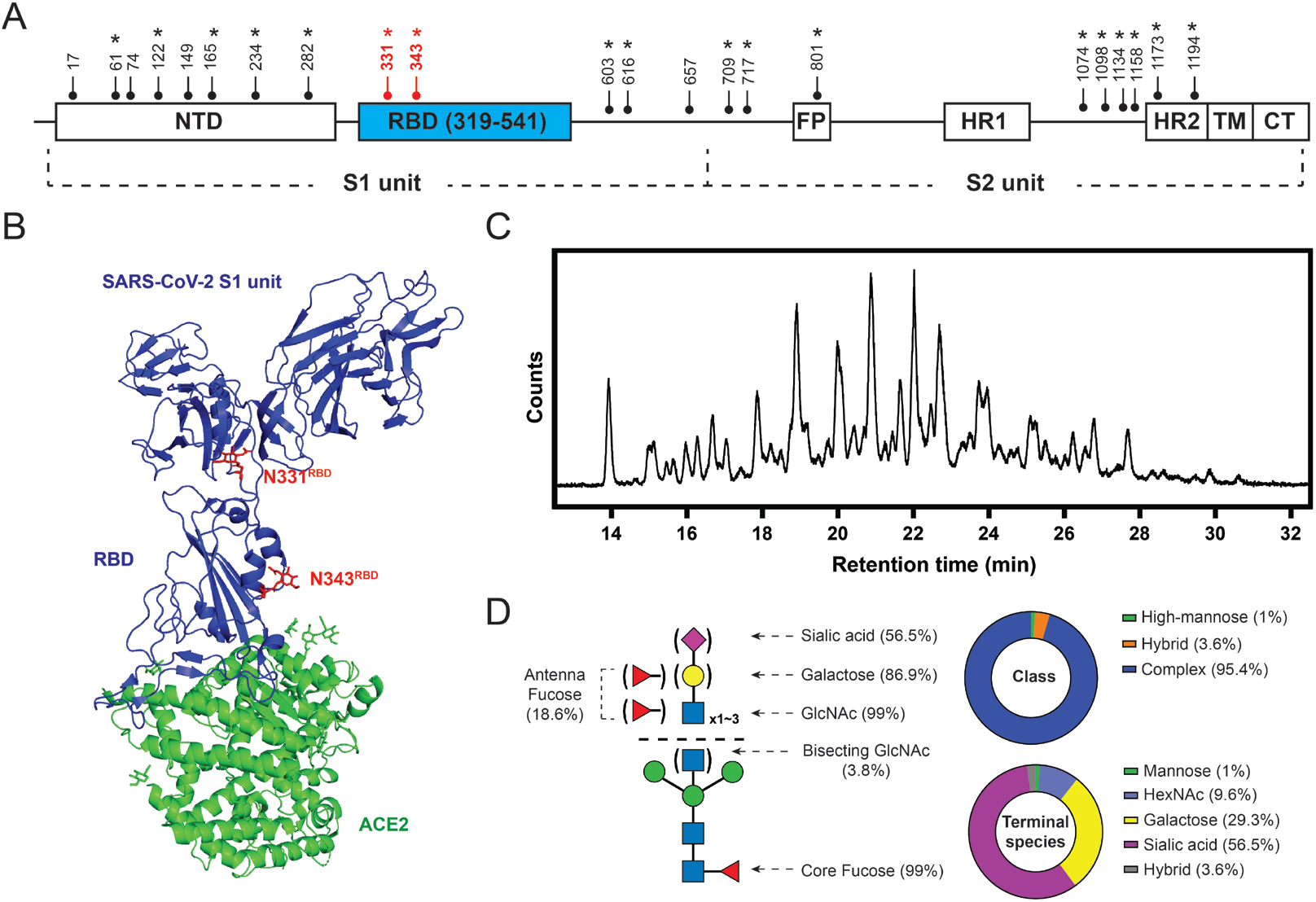
SARS-CoV-2 S protein glycosylation. (A) Glycosylation sites are distributed throughout the S protein structures. Star symbols (*) indicate conserved glycosylation sites with SARS-CoV S protein. NTD: N-terminal domain; RBD: receptor-binding domain; FP: fusion peptide; HR: heptad repeat; TM: transmembrane region; CT: cytoplasmic domain fusion. (B) Structure of the S protein S1-ACE2 complex with highlighted RBD glycosylation sites in red (Protein Data Bank (PDB): 7a92). (C) HILIC chromatogram of glycans collected from HEK293-expressed S protein RBD used in this study. (D) Composition of S protein RBD glycosylation described in relative abundance of glycoform(s) with each property. Left: graphic representative of the RBD glycopattern. Upper right: composition of glycan classes. Bottom right: composition of terminal saccharide species. 56.5% of the RBD glycan population contains at least one terminal sialic acid; 29.3% of the population exhibits at least one terminal galactose without having any sialic acid; 9.6% of the population terminating with GlcNAc without having any galactose and sialic acid.

Glycosylation plays versatile roles in viral pathobiology.(10) For example, glycans are part of the cell entry machinery of human immunodeficiency virus 1 (HIV-1). Deleting certain glycosylation sites from HIV-1 envelope protein reduces virus integrity and results in significantly decreased infectivity.(11) Similar phenomenon in SARS-CoV-2 was reported recently by Casalino *et al*., where they found that the N165 and N234 glycans on the S protein S1 subunit modulate the protein conformational transition required for receptor binding.(12) Removal of these two glycosylation sites through mutation leads to 10-40% reduction of ACE2 binding response, as supported by biolayer interferometry (BLI) analyses.(12) Another well-known role of viral glycosylation is masking viral immunogenic epitopes from the host immune systems by camouflaging them with host-derived glycans.(13, 14) This has proved to be a powerful strategy to maintain the infectivity in coronavirus S protein, HIV-1 envelope protein, and influenza hemagglutinin.(15–19) In SARS-CoV-2, the surface of the S protein is extensively covered by glycans, indicating effective camouflaging effects against antibody recognition. However, a notable exception was found in the receptor-binding domain (RBD).(9)

The S protein RBD contains 222 residues (R319 to F541 residues) with two glycosylation sites located at N331 and N343. Strong ACE2 interactions were found in the receptor-binding motif (S438 to G504 residues) through hydrogen bonds and salt bridges, as revealed by crystal structures.(20, 21) **(Figure 1B)** Despite its crucial role in ACE2 binding, the RBD has the lowest coverage of glycan shielding among the entire protein, which makes it vulnerable to immune recognization.(12, 13) One possible explanation to this phenomenon is the existence of the “up and down” conformational change of the spike protein during the cell entry processes, where the RBD remains buried by the heavily glycosylated S1 unit (the “down” state) during trafficking for immune evasion until it engages ACE2 at the infection interface (turning into the “up” state).(3, 22) This strategy minimizes the exposure time of RBD to the surrounding local environment, reducing the probability of immune recognition. However, this explanation makes the roles of the RBD glycans even more intriguing, especially for the glycans at N343 that is very close to the receptor-binding motif (RBM).(20) We suspect that the RBD glycans could have roles additional to glycan shielding in the S protein-ACE2 interactions.

In-depth probing of glycans’ functions during viral infection is challenging, largely due to the lack of strategies to control glycan structures and minimize their micro-heterogeneity.(23–26) As a result, molecular dynamics (MD) simulation has become the predominant approach for studying glycan biochemistry; yet support from experimental data is in great demand.(27) In this work, we report the strategies to harmonize the glycans of SARS-CoV-2 S protein RBD into controlled glycoforms, as well as subsequent binding affinity measurement between human ACE2 and the glycoengineered RBD. By the designed combinations of glycoengineering enzymes that we have characterized, we successfully transformed the S protein RBD glycans into 1) glycoforms with harmonized terminal glycan species; and 2) structure-defined single glycoforms.(28) This work reveals the regulatory roles of S protein RBD glycan in receptor binding: a double-edged sword that can either stabilize or destabilize RBD-ACE2 interactions. In combination with their roles in glycan shielding, these insights lay the foundations for modulating the S protein’s nature through glycan remodeling, which may create new strategies for vaccine and therapeutic design.

## Results

### Dissecting the glycoforms of SARS-CoV-2 S protein RBD

Glycans in the S protein RBD comprise N-glycan species.(6, 7) Their biosynthesis occurs in the endoplasmic reticulum (ER) and continues in the Golgi in a species-, cell-, protein-, and site-specific manner. N-glycans share a common core structure made of two *N*-Acetylglucosamine (GlcNAc) and three mannose (Man) residues. The core further extends into a myriad of glycoforms through the activity of glycosidases and glycosyltransferases. It has been known that different glycoforms could lead to different regulatory effects to protein-protein or cell-cell interactions.(29)

Given that glycan formation is sensitive to the expression conditions, we first analyzed the glycoforms of the recombinant RBD (expressed from HEK293, GenScript Z03483) used in this study by isolating the glycans using PNGase F and profiling them by liquid-chromatography mass spectrometry (LC-MS) analysis. (**Figure 1C**) Our substrate RBD exhibits heterogeneous glycoforms composed of complex-type species (95.4%), with three antennae the highest number we observed, and small amount of high-mannose (1%) and hybrid-type (3.6%) species. **(SI-Data)** As summarized in **Figure 1D**, GlcNAc, galactose, and sialic acid (SA) appear as the terminal monosaccharide at the non-reducing ends of N-glycans with relative abundances of 9.6%, 29.3%, and 56.5%, respectively. Over 99% of the RBD glycans contain the core fucose and 18.6% of the glycoforms have additional fucose located at the complex-type glycan antennae, as supported by glycosidase treatment experiments. **(Figure S1)** *N*-Acetylgalactosamine (GalNAc), the epimer of GlcNAc, likely exists in the RBD glycans since *N*-Acetylglucosaminidase (GlcNAcase) alone was not able to remove all the *N*-acetylhexosamine (HexNAc) residues, unless GalNAcase was also added. **(Figure S2)** In addition, retention time comparison using glycan standards suggested that a small amount of bisecting N-glycan species exist, as indicated by FA2B glycoform ([Man]_3_[GlcNAc]_5_[Fuc]_1_). **(Figure S3)** We note that our analyses based on isolated glycans did not provide site-specific information for the glycosidic linkages. The reported numbers here are averaged results from the glycan populations at N331 and N343.

### The RBD glycans can stabilize RBD-ACE2 interaction

We used BLI to study the interaction between the S protein RBD and ACE2 expressed from HEK293. The RBD with native glycoforms binds to ACE2 with an equilibrium dissociation constant (*K_D_*) of 99 ± 12 nM, similar to earlier reports.(30) The terminal saccharides of mature glycans often regulate the biochemical properties and functions of glycoconjugates. A good example comes from the human blood group antigens that are classified by their terminal residue species.(31) Therefore, we aimed to harmonize the RBD glycan termini and investigate their potential effect on the RBD-ACE2 interaction. First, we created glycoengineered RBD glycoprotein with all N-glycans ending in terminal HexNAc by incubating the native RBD with α2-3/6/8 neuraminidase, β1-4 galactosidase, and α1-2,3/4 fucosidases. **(Figure 2, Table S1)** Interestingly, we found that the resulting substrate, named tHexNAc-RBD, had improved binding affinity to ACE2 with a *K_D_* value of 47 ± 8 nM. **(Figure 3A-B)** Direct comparison of ACE2 binding curves showed a 20% increase in binding response in tHexNAc-RBD compared to the native RBD. **(Figure S4)** Furthermore, we prepared glycoengineered RBD bearing (i) the core glycan ([Man]_3_[GlcNAc]_2_[Fuc]_1_, core-RBD), and (ii) species terminating with galactose (tGal-RBD) by introducing *N*-Acetylglucosaminidase/*N*-Acetylhexosaminidase cocktail and Galactosyltransferase to the reactions, respectively. Similar to the tHexNAc-RBD, having glycans terminating with mannose and galactose slightly improved the ACE2 binding affinity relative to the native RBD. This result suggests that certain glycoforms can facilitate RBD-ACE2 interactions.

**Figure 2.**
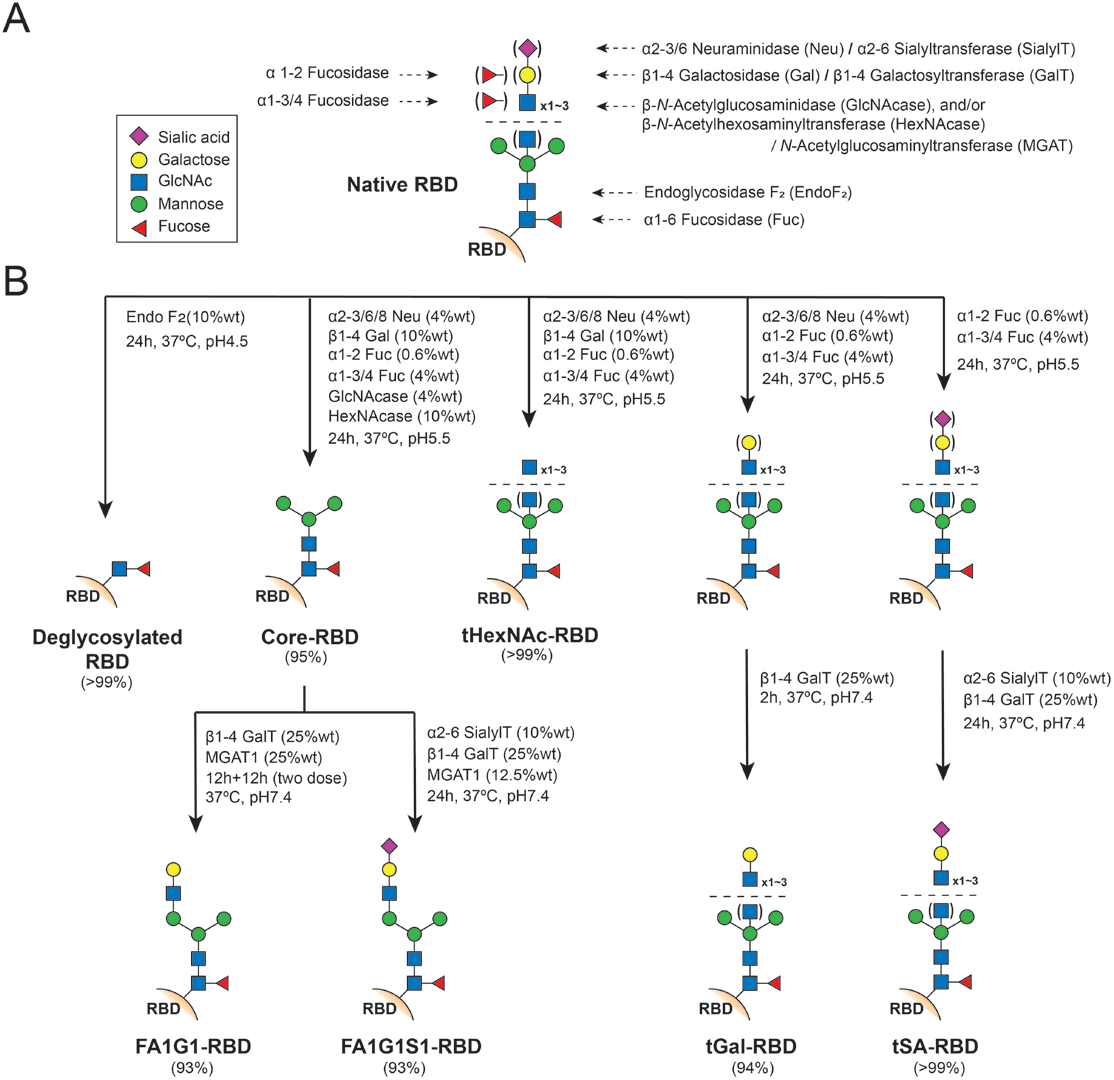
Scheme of the S protein RBD glycan remodeling routes. (A) Enzymes used in this work and their corresponding saccharide targets. (B) Preparation of the S protein RBD with controlled glycoforms using enzymatic reactions. The yield indicates estimated populations of the desired glycoform(s).

**Figure 3.**
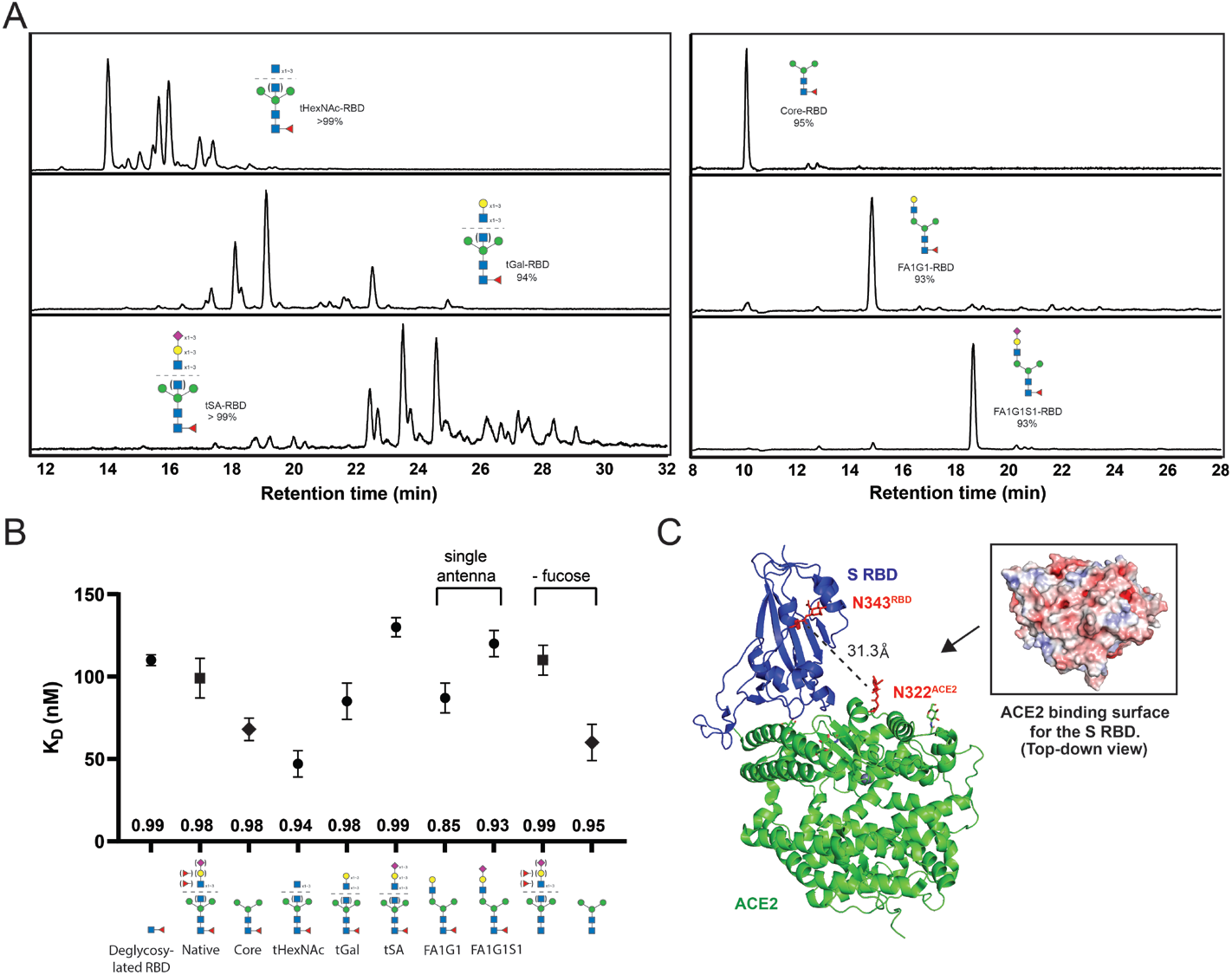
Glycans of the S protein RBD affect ACE2 binding affinity. (A) Chromatograms of glycans collected from glycoengineered RBD in this study. Left: the RBD with harmonized terminal glycan species. Right: the RBD substrates with single mono-antennary glycoforms. (B) Binding affinity measurement (BLI) between human ACE2 and the S protein RBD with controlled glycoforms. The numbers located at the bottom of each data point indicate the R^2^ value of curve fitting. Error bars: standard error of mean. (C) Structure of the RBD-ACE2 complex revealed potential interaction between the RBD N343 glycan and ACE2 glycans/amino acid backbone (PDB: 6m0j). Insertion displays the electrostatic potential of ACE2 surface that potentially interacts with RBD N343 glycan. Red: negatively charged; blue: positively charged.

Recently published simulation studies by Mehdipour *et al*. provided a plausible explanation to our observation.(32) In addition to the S protein RBD, ACE2 is also a heavily glycosylated protein.(33) Both the RBD glycan at N343 (N343^RBD^) and the ACE2 glycan at N322 (N322^ACE2^) are close to the binding interface with a distance of only 31.3 Å between the two glycans (Asn side chains, **Figure 3C**). Mehdipour and co-workers investigated the ACE2 glycan dynamics and found that the N322^ACE2^ glycans interact with the RBD protein backbone as well as the N343^RBD^ glycans. Modeling analysis indicated that these interactions generate 250-360 kJ/mol binding force, which contributes to about one-third of the RBD-ACE2 binding energy (890-1040 kJ/mol).(32) In other words, the RBD glycans interact with ACE2, which explains why remodeling their structures led to different binding affinities. To examine this explanation, we measured the ACE2 binding affinity of deglycosylated RBD prepared by endoglycosidase F2 treatment.(34, 35) **(Figure S5)** An apparent reduction in binding response was found with a *K_D_* value of 110 ± 3.3 nM, a weaker affinity than that of native RBD. **(Figure 3B)** Moreover, when the ACE2 N-glycans were removed, a dramatic decrease of binding affinity was observed. **(Figure S6)** In combination with the published simulation study, our result suggested that the RBD glycans can stabilize ACE2 binding, likely through the interactions with the N322^ACE2^ glycans or the ACE2 backbone.

### Glycans with terminal sialic acid destabilize RBD-ACE2 interactions

Sialic acids are often involved in biomolecular interactions because they are usually found at glycan non-reducing end termini. For example, binding to sialic acid and subsequent release by neuraminidase are required events for influenza virus infection.(36) By incubating α2-6 sialyltransferase with the tGal-RBD, we introduced sialic acids to the terminus of the RBD glycans. The resulting tSA-RBD contains heterogeneous glycoforms with over 99% of the population bearing at least one terminal sialic acid. **(Figure 3A, SI-Data)** Unexpectedly, BLI measurement showed that the tSA-RBD has lower binding affinity to ACE2 (*K_D_* = 130 ± 6 nM) compared to the Native and deglycosylated RBD. We reasoned this result could be attributed to the electrostatic repulsion between the N343^RBD^ glycan and the ACE2 surface. Computational calculation of electrostatic potential has shown that the ACE2 surface is predominantly negative, including the area that N343^RBD^ glycan engages.(37) **(Figure 3C)** Therefore, sialylated glycoforms, which are negatively charged, could cause electrostatic repulsion and destabilize RBD-ACE2 interactions. Given that the native RBD has a heterogeneous glycan profile, the existence of the sialylated glycan species possibly negates the stabilizing effects resulting from other glycoforms, which explains the weaker ACE2 binding affinity in native RBD over non-sialylated glycoforms. Recent reports of site-specific glycan mapping have also indicated that both native N343^RBD^ and N322^ACE2^ glycosylation sites have a low content of sialylated glycan species, further implying that sialylation may not be favored for RBD-ACE2 interactions.(7, 33)

Another possible explanation of the reduced ACE2 binding affinity found in tSA-RBD is the increased steric hindrance caused by sialylation. Having over-constructed glycan structures nearby the receptor-binding motif could prevent the S protein from approaching ACE2. To test this possibility, we constructed harmonized mono-antennary glycans on RBD, as confirmed by LC-MS analysis **(Figure 3A, S7, SI-Data)**. The FA1G1-RBD bears an octa-saccharide with only one antenna connected to the α1-3-linked core mannose. This was achieved by incubating the core-RBD with *N*-acetylglucosaminyltransferase I (also known as MGAT1) and galactosyltransferase in one pot. Similarly, the FA1G1S1-RBD contains an additional sialic acid and was prepared by introducing sialyltransferase to the reaction. BLI analyses showed that the FA1G1-RBD has improved ACE2 binding affinity compared to native RBD; by contrast, FA1G1S1-RBD showed weaker binding than native RBD, which is consistent with the data found earlier. **(Figure 3B)** This result suggested that steric hindrance is not the major cause of the ACE2-binding affinity reduction found in the tSA-RBD. Instead, the electrostatic repulsion is more likely the reason. Our results, together, suggest that the RBD glycans have regulatory effects on the RBD-ACE2 interactions: non-sialylated glycoforms reinforce the binding while sialylated glycoforms destabilize it.

### Fucose in the RBD glycans does not have apparent impacts on ACE2 binding

Glycosidase treatment provided direct evidence that the core structure of the complex-type RBD glycans is fucosylated because afucosylated core was not detected. **(Figure 3A, S1)** The core fucose linked to the reducing end GlcNAc has been a hot target of interest for pharmaceutical research because of its regulatory effects on protein-protein interactions, such as immunoglobulin G (IgG) and its receptors.(38) To investigate the role of fucosylation in the RBD-ACE2 interaction, we prepared partially defucosylated RBD using α1-6 fucosidase to target the core fucose. A 10-day reaction converted approximately 45% native glycans into non-fucosylated forms. **(Figure S8)** It is known that the activity of this enzyme is glycoform-dependent, where lower structural complexity leads to higher enzyme activity.(28) Therefore, we used the core-RBD as the substrate for the reaction and successfully remove fucose from over 90% of the core glycans. **(Figure S8)** BLI measurement showed that both defucosylated RBD substrates have no significant difference in ACE2 binding affinity compared to their parent species. **(Figure 3B)** This result is not surprising because the core fucose is shielded close to the RBD peptide backbone, which minimizes its probability of interacting with ACE2 or ACE2 glycans. In addition to the core-fucose, approximately 18.6% RBD glycans contain α1-2 and α1-3/4 fucose which can be removed by corresponding fucosidases. **(Figure S1)**. Their removal did not result in a significant difference in ACE2 binding affinity (*K_D_* = 110 ± 14 nM).

### Monoclonal Antibody S309 neutralizes the regulatory effects of the RBD glycans

Neutralizing antibodies have presented effective therapeutics for viral infection treatment.(39) An antibody, named S309, isolated from a SARS-CoV-infected patient was reported to have strong cross-neutralization activity on SARS-CoV-2 through the induction of antibody-dependent cell cytotoxicity (ADCC) and antibody-dependent cellular phagocytosis (ADCP).(40) S309 has a distinct RBD binding interface from that of ACE2. It recognizes a SARS-CoV-2 S protein RBD epitope consisting of two regions, residues 356-361 and residues 440-444. Notably, the S309 binding interface sandwiches the N343^RBD^ glycosylation site and shows potential interaction with the glycan structure, especially the core fucose.(40) **(Figure 4A)** To investigate this interaction, we measured the binding affinity between S309 and the RBD with differently engineered glycan structures. BLI analyses showed strong RBD-S309 interaction with *K_D_* values at nanomolar scale. However, no significant difference was found when the RBD glycan bore different terminal saccharide species, so as the removal of the core fucose. **(Figure 4B)** This result suggested that the N343^RBD^ glycan is not essential for the RBD-S309 binding. The RBD-S309 interaction is likely established on the amino acid epitope of the RBD.

**Figure 4.**
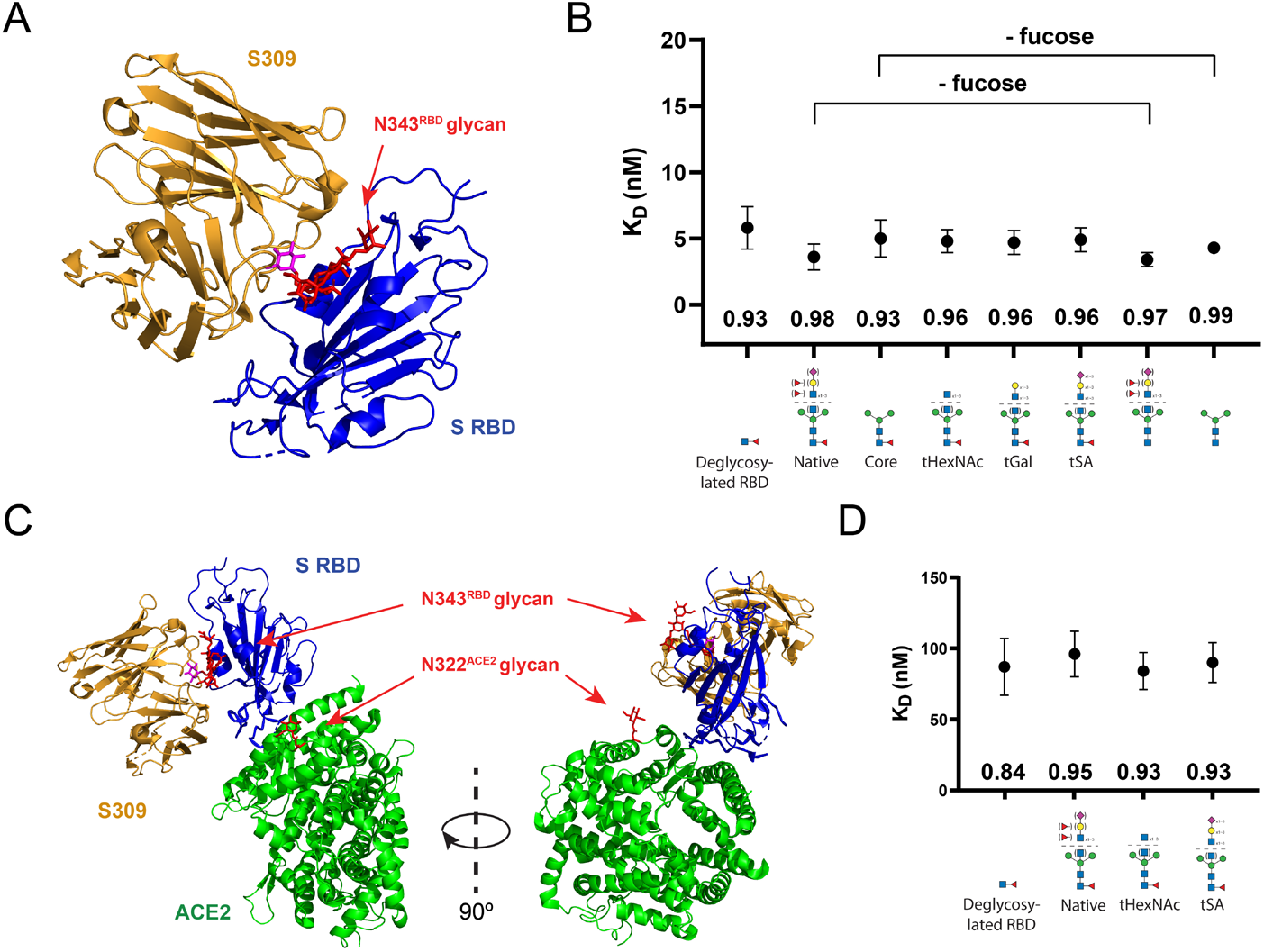
Antibody S309 neutralizes the regulatory effect of the RBD glycans. (A) The N343^RBD^ glycan is located at the RBD-S309 binding interface. (PDB: 6wps) The core fucose is highlighted in pink. (B) RBD glycans are not involved in S309 binding. No significant difference was observed in the binding affinity analysis. (C) The RBD-S309 interaction hinders the accessibility of the N343^RBD^ glycan to ACE2 and ACE2 glycans. The composite model was generated using the RBD-S309 cryo-EM structure (PDB: 6wps) reported by Pinto *et al*. and the RBD-ACE2 crystal structure (PDB: 6m0j) reported by Lan *et al*.^20, 39^ (D) The RBD-S309 binding neutralizes the regulatory effect of the RBD glycan on ACE2 interaction. The numbers located at the bottom of each data point indicate the R^2^ value of curve fitting. Error bars: standard error of mean.

Despite the N343^RBD^ glycan is not involved in the antibody binding, its structure has been sterically hindered by the S309-RBD interaction, which could restrict its probability of interacting with ACE2 and/or ACE2 glycans. **(Figure 4C)** To test this possibility, we measured the ACE2 affinity of the RBD samples that were bound to S309. As expected, comparable binding affinities (*K_D_* ≈ 90 nM) were observed on the native RBD, tHexNAc-RBD and tSA-RBD that were saturated with S309. **(Figure 4D)** Namely, S309 neutralizes the regulatory effect of the RBD glycan. Given that the native RBD glycans tend to stabilize ACE2 interaction owing to the low content of sialylated glycoforms, the RBD-S309 binding could compromise this stabilizing effect and reduce the infectivity of the virus, which presents a new inhibitory mechanism in S309 neutralization.

## Discussions

The COVID-19 pandemic has caused numerous deaths worldwide. While the emerging vaccines have partially relieved the immediate suffering, chronic symptoms could have persisted in many who have contracted the virus.(41) With the rapid evolution of SARS-CoV-2, a comprehensive understanding of the virus pathology is in urgent demand for developing vaccines and effective therapeutics to cease the spread of the virus variants.(42–44)

Glycans play versatile structural and functional roles in protein biochemistry, but it has remained historically challenging to elucidate the phenotypic roles of discrete glycoforms. Given the proximity of the RBD glycans to the binding interface with ACE2, investigating their roles in its biological properties could provide insights into the infection mechanism.(7, 8) We have used an enzymatic remodeling approach to remove the native glycosylation microheterogeneity and decorate the S protein RBD with specific glycan structures in order to study their impact on receptor binding.(28) We found that the neutrally charged RBD glycans can stabilize ACE2 binding, likely through the interaction with ACE2 glycans, as suggested by published simulation studies.(32) Remarkably, this effect is reversed when negatively charged sialic acids are present on the RBD glycans, a destabilizing force caused by electrostatic repulsion. The dissociation constants measured herein varied approximately 3-fold, based solely on the displayed N-glycan structures. The results suggest that SARS-CoV-2 infectivity could be altered by manipulating the content of sialic acid on the RBD glycans. For example, neuraminidase inhibitors could mitigate the viral infection by halting the removal of sialic acid from the spike protein. Although anti-influenza neuraminidase inhibitors have proven to be inactive for SARS-CoV-2 treatment, strategies that regulate the neuraminidase and sialyltransferase activities on the RBD glycans are still worth investigating.(45)

In addition, we found that the binding of S309 neutralizes the regulatory effects of the RBD glycan, preventing the potential stabilization of ACE2 interaction through the glycan structure. Given that the glycosylation sites of the RBD are highly conserved across its evolution reported to date, glycans could provide new handles for developing strategies to combat the emerging virus variants **(Figure S9).** For example, glycan remodeling could be applied to protein subunit vaccines to maximize the mimicry of viral glycoproteins and improve the immune response. Generally speaking, this work demonstrates that it is possible to efficiently screen the biological behavior of discrete protein glycoforms, an approach that will greatly advance our fundamental knowledge of glycobiology and reinforce our ability to design targeted strategies to defeat disease.

## Supporting information

Supplementary Data (LCMS)

## Data Availability

All data are available upon reasonable request to the corresponding authors.

## Author contributions

Method development for glycan remodeling, binding affinity measurements, LC-MS analysis, and data analysis was performed by Y-P. H. MALDI-ToF analysis was performed by D. M. and V. S. All the authors were involved in the design of the research and manuscript drafting.

## Acknowledgment

We thank the Merck Research Laboratory (MRL), Merck Postdoctoral Research Fellows Program, and Analytical Research and Development (AR&D) department for the financial support. We are also thankful to Erik Regalado and Jimmy DaSilva (Method Screening and Purification group) for technical assistance and advice; Jeffrey Moore and David Row (Protein Engineering group) for kind suggestions on the experimental designs. We also thank Caroline McGregor and Ian Mangion for their support of this study in the AR&D department.

