## Supplementary Materials for "Structural remodeling of SARS-CoV-2 spike protein glycans reveals the regulatory roles in receptor binding affinity"

Raw data that support the findings of this study are available upon reasonable request.

Proprietary

**Materials and Instruments**

SARS-CoV-2 spike protein receptor-binding domain (Arg319-Ser591, with an Avi/His tag at the C-terminal) was purchased from GenScirpt (Z03483). Recombinant human ACE2 (10108-H02H) was purchased from SinoBiological. Glycosidases were purchased from NEB: α2-3/6/8 Neuraminidase (P0720), β1-4 Galactosidase (P0745), β-N-Acetylglucosaminidase (P0744), β-N-Acetylhexosaminidase (P0721), α1-2 Fucosidase (P0724), α1-3/4 Fucosidase (P0769), α1-6 Fucosidase (P0749), Endoglycosidase H (P0702), Endoglycosidase F2 (P0772). Glycosyltransferases were purchased from Sigma-Aldrich and R&D Systems: α2-6 Sialyltransferase (SialylT, SAE0090), β1-4 Galactosyltransferase (GalT, SAE0093), β1-2 N-Acetylglucosaminyltransferase (MGAT1, 8334-GT). Monoclonal antibody S309 was produced by Cell Sciences.

Ni-NTA resin (88221) was obtained from ThermoFisher. Tris-HCl (T-5941), HEPES (H4034), sodium acetate (S2889) sodium chloride (C5670), magnesium chloride (M4880), manganese chloride solution (M1787), sodium chloride solution (S5150), CMP-NANA (C8271), UDP-Gal (U4500), UDP-GlcNAc (U4375), Acetonitrile (900667), Discovery glycan SPE columns (55465-U), and empty SPE columns and frits (57607-U) were purchased from Sigma-Aldrich. HILIC columns for chromatography (186004742), SPE µPlate (186002780), µPlate extraction manifold (186001831), SPE vacuum manifold (WAT200607), RapiGest SF (186008090), Glycoworks buffer (186008100), glycan standards (186008791), and Rapifluor-MS (186008091) were purchased from Waters. 10K MWCO filters (UFC503096) were purchased from Millipore Sigma. Rapid PNGaseF (P0710) was purchased from NEB. Octet sensors (18-5101) were purchased from Sartorius.

Chromatography and mass spectrometry analyses were conducted using an Agilent 1290 Infinity II LC system tandem with Agilent 6500 Series quadrupole time-of-flight (TOF) MS system. Enzyme concentration was determined by absorbance at 280 nm using NanoDrop 2000 (Thermo Scientific). pH value was measured by using a Mettler Toledo S220 pH meter. Temperature-controlled reactions/incubations were performed in ThermoFisher MaxQ 6000 incubator and Fisherbrand Thermal Mizer II. Biolayer interferometry analysis was performed by Octet$^{Red}$ 384 system (ForteBio, now Sartorius).

**Methods**

The Spike protein RBD glycan remodeling

Please see **Figure 2 and Figure S1** for reaction conditions and remodeling routes. Reactions were started by mixing the S protein RBD and glycoengineering enzyme(s) in a 10K MWCO tube for buffer exchange into the corresponding reaction buffer. The reaction solution was then collected into a 1.7 ml microtube and incubated in a temperature-controlled shaker until the reaction was complete, as confirmed by glycan analysis (see below). Standard Ni-NTA resin purification protocol was used to purify the crude product (empty SPE column and vacuum manifold were used for the purification). Finally, the purified product was transferred into 10 K MWCO tubes for buffer exchange into either Octet buffer (25 mM Tris-HCl pH 7.4, 50 mM NaCl, 0.02% Tween 20) for biolayer interferometry analysis or the reaction buffer for the subsequent reaction. The enzyme and RBD concentrations were determined by NanoDrop. The glycan remodeling processes can be simplified by combining certain enzymes into one-pot reactions. Please refer to **Figure 2** for more details.

Pr Proprietary

The reaction buffer was prepared as follows: For glycosidase reactions: 50 mM sodium acetate at pH 5.5 with 5 mM $CaCl_2$. For glycosyltransferase reactions: 25 mM Tris-HCl pH 7.4 with 50 mM NaCl and 10 mM cations ($MnCl_2$, $MgCl_2$, $CaCl_2$). The buffer for Endoglycosidase reactions contains 50 mM sodium acetate at pH 4.5 for Endo $F_2$ and Endo $F_3$; 50 mM sodium acetate at pH6 for Endo H and Endo M; 50 mM sodium acetate at pH 5.5 and 5 mM $CaCl_2$ for Endo S.

LC-MS Analysis of S protein RBD glycans

This protocol is adapted from the Glycoworks manual provided by Waters. **(I) Glycan isolation:** 15 µg of the S protein RBD (in either reaction buffer or octet buffer), 6 µl RapiGest SF (50 mg/ml, in Glycoworks buffer), and water were mixed in a 1.5 ml microtube to a final volume of 25 µL. The mixture was then incubated at 90 °C for 5 minutes to denature the substrates. After the samples were cooled down to room temperature, 1.2 µL Rapid PNGase F was added to the tube, followed by another incubation at 50 °C for 10 minutes. **(II) Glycan labeling:** After the PNGase F digestion, 12 µL RapiFluor-MS solution (70 mg/ml, in DMF) was added directly to the solution. The mixture was gently vortexed and then incubated at room temperature for 20 minutes without any light exposure. RapiFluor-MS labeling gives higher signal-to-noise (S/N) ratio in MS analysis. A larger amount of analyte might be required without the labeling step. After labeling, the samples (~40 µL) were diluted with 360 µL acetonitrile and were ready for purification. **(III) Glycan purification:** Oasis SPE µPlate from Waters (along with the use of µPlate extraction manifold) was then used for the 1st solid-phase extraction purification, and Discovery SPE from Sigma-Aldrich (along with the use of 20-wells SPE vacuum manifold) was used for the 2nd purification to ensure high signal-to-noise ratio. The SPE columns/µPlate were first washed by water (1 column volume) and then conditioned by water-acetonitrile solution (10:90 v/v, 1 column volume). The glycan samples (in ~90% acetonitrile solution) were then charged to the column/µPlate, followed by washing with washing buffer (formic acid/water/acetonitrile 1:9:90 v/v/v, 2 column volume). The glycans were then eluted with 80 µL elution buffer (200 mM ammonium acetate in 5% acetonitrile). **(IV) HILIC-MS analysis:** The purified glycan samples were injected to UPLC equipped with ACQUITY BEH Glycan column (130 Å, 1.7 µm, 2.1 x 150 mm) tandem with IMQ-TOF MS for glycan profile analysis. The method provided by Waters for glycan chromatography was used in this work:

**Mobile phase A:** 50 mM ammonium formate in $H_2O$, pH 4.4
**Mobile phase B:** 100% Acetonitrile
**Temperature:** 60 °C
**Injection volume:** 5-10 µL

| Time (min) | Flow rate (ml/min) | %A | %B |
|---|---|---|---|
| 0 | 0.4 | 25 | 75 |
| 35 | 0.4 | 46 | 54 |
| 36.5 | 0.2 | 100 | 0 |
| 39.5 | 0.2 | 100 | 0 |
| 43.1 | 0.2 | 25 | 75 |
| 47.6 | 0.4 | 25 | 75 |
| 55 | 0.4 | 25 | 75 |

The instrument setting for the collection of mass spectrometry data was as follows: Source type: Dual AJS ESI with gas temperature at 300 °C; Nebulizer pressure at 35 PSIG; Sheath gas Temperature at 325°C; Nozzle voltage at 2kV; and Fragmentor voltage: 175V. The scan range was 600−3200 m/z. Only intact mass data was collected. System tuning was performed using Agilent tune mix (G1969-8501, G1969-8503)

Glycan analyses

The N-glycans on the S protein RBD were determined by LC-MS analysis. MassHunter Software from Agilent was used to acquire and analyze the data. **(I) Peak area integration:** the area of each chromatographic peak was determined by drawing the peak baseline manually in order to include the glycoforms with low abundance. Peaks with an S/N ratio less than 1.2 were excluded. **(II) Glycan assignment:** the collected MS data from each peak was imported into the GlycanMass and GlycoMod analysis tools operated by Expasy to reveal the saccharide composition of the glycans.(1, 2) Given that the majority of RBD glycans belongs to complex-type species, glycoforms can be predicted by matching the saccharide compositions to the complex-type glycan database (GlycoMod). Glycan standards (Waters) were used to confirm certain glycoforms, e.g. FA2 glycan, based on retention time comparison, in addition to the accurate mass measurement. **(III) Glycoform population:** after peak assignment, the relative abundance of glycoform(s) of interest was calculated based on the UV peak area. Detailed glycan structure characterization using NMR or MS fragmentation is beyond the scope of this study and was not included.

Binding affinity analysis using biolayer interferometry (Octet)

Ni-NTA Octet sensors were used for the measurement of the S protein RBD-ACE2 binding affinity. The sensors were hydrated in Octet buffer (25 mM Tris-HCl pH 7.4, 50 mM NaCl, 0.02% Tween 20) for 30 minutes before the experiments. To begin the measurement, the sensors were loaded with 2 µg/ml (50 nM) His-tagged S protein RBD for 400 seconds, followed by baseline equilibration for 10 minutes. Association of recombinant human ACE2 was performed at 0, 10, 20, 30, 40, 60, 80, and 120 µg/ml (0-1 µM) for 400 seconds. Dissociation was measured for 30 minutes. The equilibrium dissociation constant ($K_D$) values were calculated using a 1:1 global fit model in the Octet data analysis software. The Octet buffer was used in all the steps. Measurement for the RBD-S309 binding affinity was carried out using a similar protocol with the antibody concentration at 0, 0.2, 0.4, 0.8, 1.6, 3.2, 6.4, and 12.8 µg/ml (0-85 nM).

The measurement of ACE2 binding affinity with the S309-bound RBD was carried out using adjusted protocols: an incubation with S309 at a high concentration (12 µg/ml) was introduced to the BLI measurement process after the Ni-NTA sensors were loaded with the RBD. The binding response was monitored in real-time until it reached a plateau (1200 seconds) to ensure S309 binding was saturated. The ACE2 association and dissociation measurements were then carried out using the same protocol for the $K_D$ calculation. All the measurements were repeated twice using separately prepared RBD substrate, buffers, and unused sensors. A consistent trend was obtained and the data set with the highest $R^2$ value was reported.

MALDI-TOF analysis of intact glycoengineered S Protein RBD

The MALDI-MS datasets for the glycoengineered RBD were acquired using a Bruker rapifleX MALDI-TOF instrument (Bruker Daltonics, Billerica, MA, UA) that was set to an average of 10000 laser shots per sample at a laser energy value of 90% with a scan range of 20-220 kDa. The instrument was calibrated prior to measurements using – 1) the Bruker Protein Standard II calibration kit (Bruker Daltonics, Billerica, MA,

UA), and 2) a 0.2 mg/mL solution of Bovine Serum Albumin (Sigma Aldrich, MO) in 50:50 acetonitrile:water. The mass errors for the calibration data were confirmed from the differences between observed and theoretical masses for each protein, and the error values were found to be <100 ppm. Sinapic acid (Sigma Aldrich, MO) dissolved in 50:50 acetonitrile:water, at a concentration of 10 mg/mL with 0.1% 3-Nitrobenzylalcohol (Sigma Aldrich, MO) to aid in ionization, was used as a MALDI matrix for both instrument calibration and sample measurement purposes. The MALDI-TOF data was collected using a linear mode of detection with either a detector gain factor of 1.9X or a higher gain value of 10X.

<u>Plotting and graphic</u>

Data plotting and curve fitting was done with GraphPad Prism 8. Figures were created by Adobe Illustrator. Protein three-dimensional (3D) structures were constructed in PyMOL with the use of the published crystal models, PDB 6m0J and 7a92.(3, 4) The protein surface electrostatic potential map was generated by the APBS Electrostatics plugin of PyMOL.(5, 6)

### Tables

| Enzyme | Source | Function (on the RBD glycans) | Class |
|---|---|---|---|
| α2-3/6/8 Neuraminidase (Sialidase) | *C. perfringens* | Terminal α2-3/6-linked sialic acid removal | Exoglycosidase |
| β1-4 Galactosidase | *S. pneumoniae* | Terminal β1-4-linked galactose removal | Exoglycosidase |
| β-*N*-Acetylglucosaminidase (GlcNAcase) | *S. pneumoniae* | Terminal β-linked GlcNAc removal | Exoglycosidase |
| β-*N*-Acetylhexosaminidase (HexNAcase) | *S. plicatus* | Terminal β-linked GlcNAc and GalNAc removal | Exoglycosidase |
| α1-2 Fucosidase | *X. manihotis* | Terminal α1-2-linked fucose removal | Exoglycosidase |
| α1-3/4 Fucosidase | *P. dulcis* | Terminal α1-3/4-linked fucose removal | Exoglycosidase |
| α1-6 Fucosidase | *C. Omnitrophica* | Terminal α1-6-linked fucose removal | Exoglycosidase |
| Endoglycosidase H | *S. plicatus* | High-mannose and hybrid-type glycans removal | Endoglycosidase |
| Endoglycosidase F2 | *E. miricola* | High-mannose and complex-type glycans removal | Endoglycosidase |
| α2-6 Sialyltransferase (SialylT) | *H. sapiens* | Conjugate sialic acid to terminal galactose through α1-6 linkage | Glycosyltransferase |
| β1-4 Galactosyltransferase (GalT) | *H. sapiens* | Conjugate galactose to terminal GlcNAc through β1-4 linkage | Glycosyltransferase |
| *N*-Acetylglucosaminyltransferase (MGAT1) | *H. sapiens* | Conjugate GlcNAc to terminal α1,3-linked mannose through β1,2 linkage | Glycosyltransferase |

**Table S1.** Glycoengineering enzymes used in this study.

Pr Proprietary

### Figures

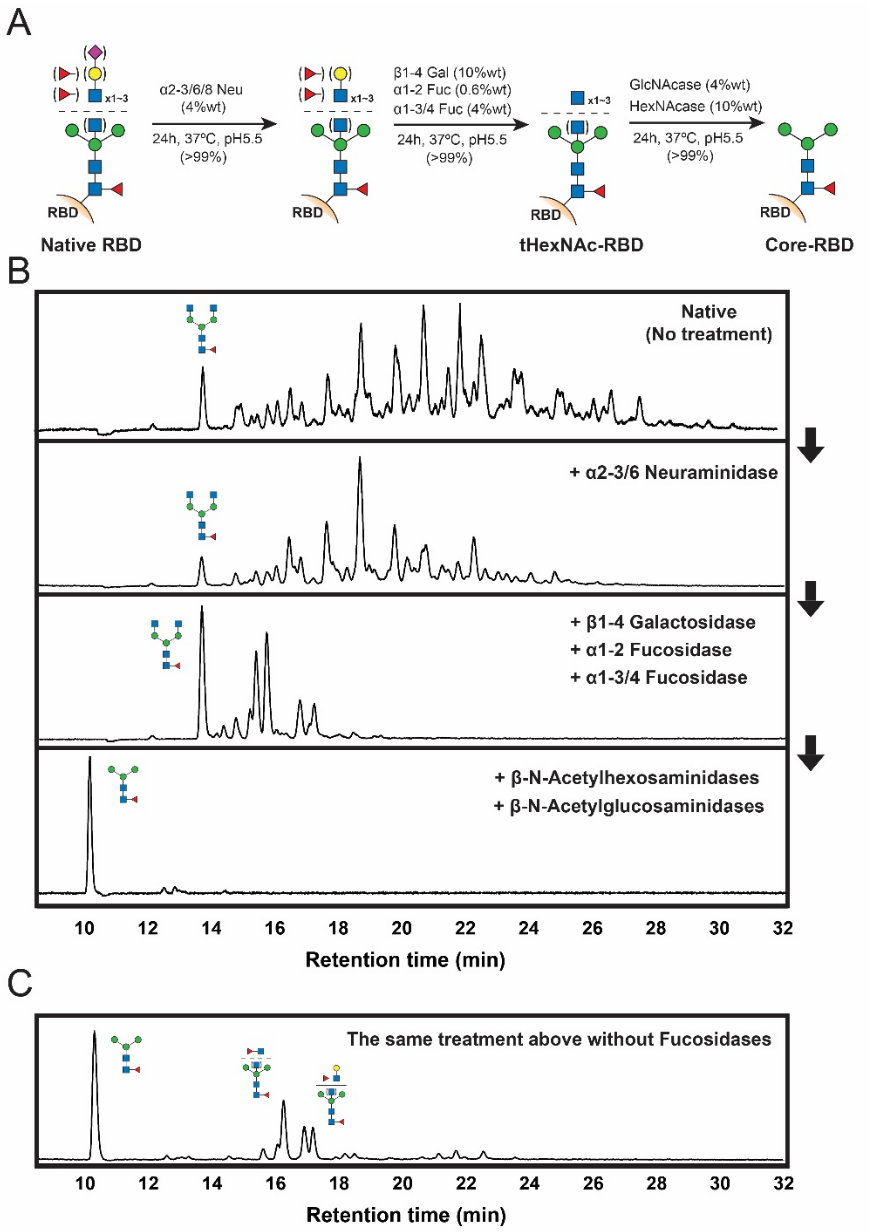


**Figure S1.** Dissecting SARS-CoV-2 S protein RBD glycoforms by sequential glycosidase treatment. (A) Scheme of the enzymatic reactions (B) Chromatograms of glycans collected from the product RBD. Sequential treatment led to the fucosylated core structure. β-N-Acetylhexosaminidase removes both terminal N-Acetylglucosaminidase (GlcNAc) and N-Acetylgalactosamine (GalNAc) (C) α1-2 and α1-3,4 fucoses are present on the RBD glycan antennae. This chromatogram was used for calculating the population of antennary fucose species.

Proprietary

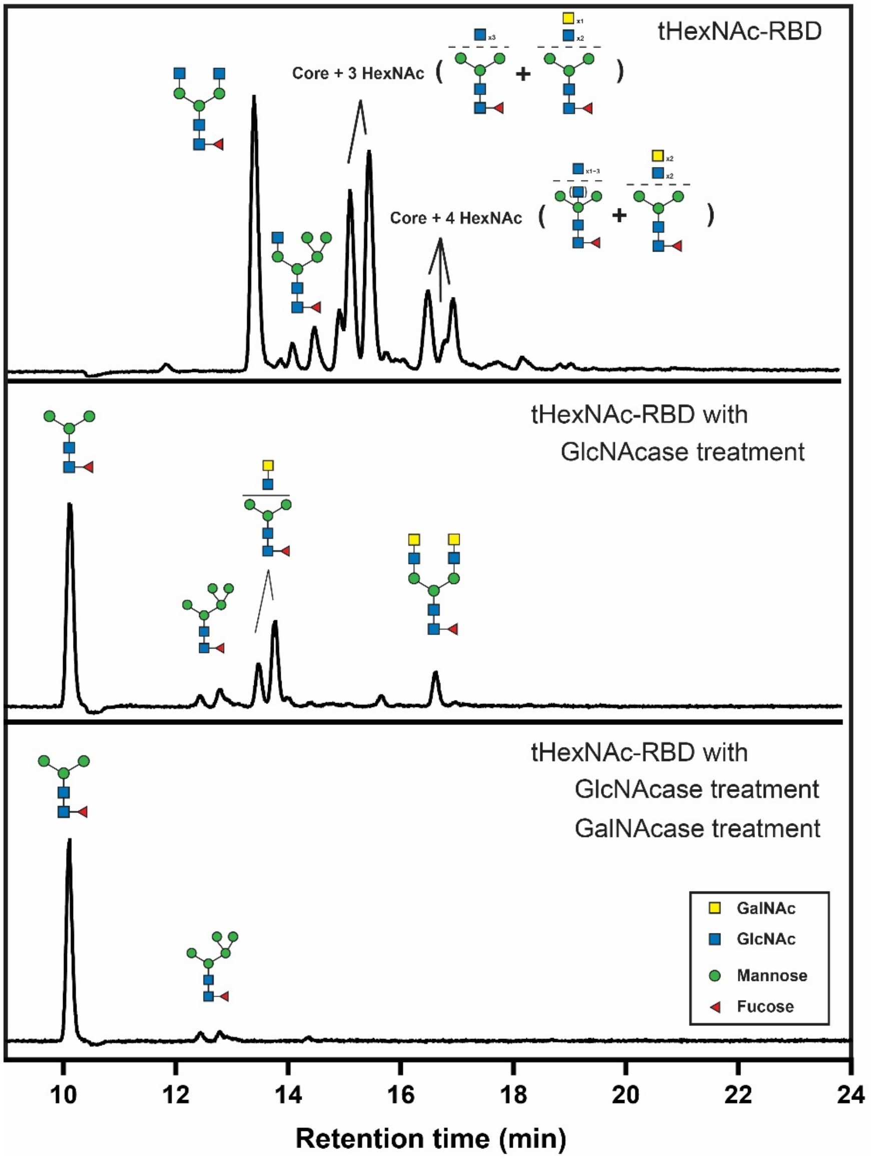


**Figure S2.** The RBD glycan contains N-Acetylglucosamine (GlcNAc) and N-Acetylgalactosamine (GalNAc). Saccharide composition analysis showed that HexNAc residues remained on the RBD glycans after the GlcNAcase treatment of tHexNAc-RBD. The residual HexNAc residues were removed by adding HexNAcase that has both GlcNAcase and GalNAcase activity. According to the analysis, approximately 30% of the RBD glycans contain at least one GalNAc.

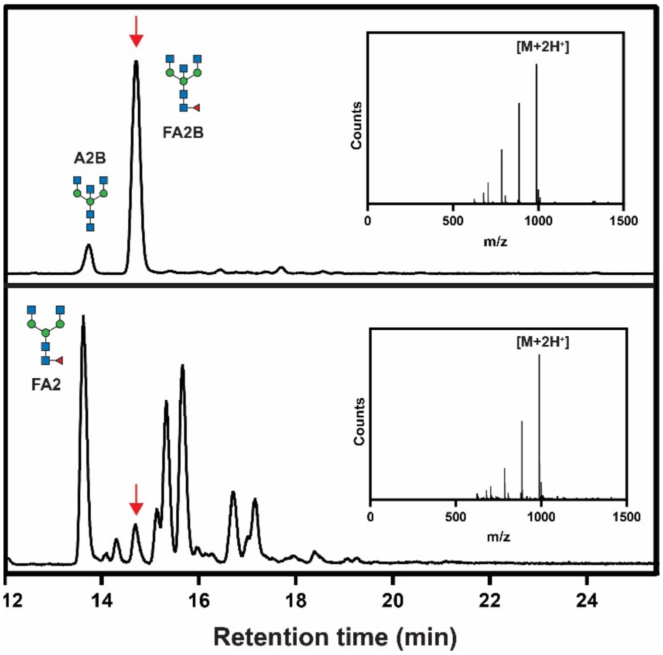


**Figure S3.** Bisecting N-glycan species were detected in the SARS-CoV-2 S protein RBD based on retention time analysis using glycan standards. Upper panel: bisecting glycan standards (A2B and FA2B, Oxford nomenclature) collected from IgG. Bottom panel: tHexNAc-RBD prepared in this work. The FA2B glycan was indicated by red arrows. Insertions are the mass-spectra of the FA2B peak. Whether or not other bisecting species exist remains to be investigated.

Pr Proprietary

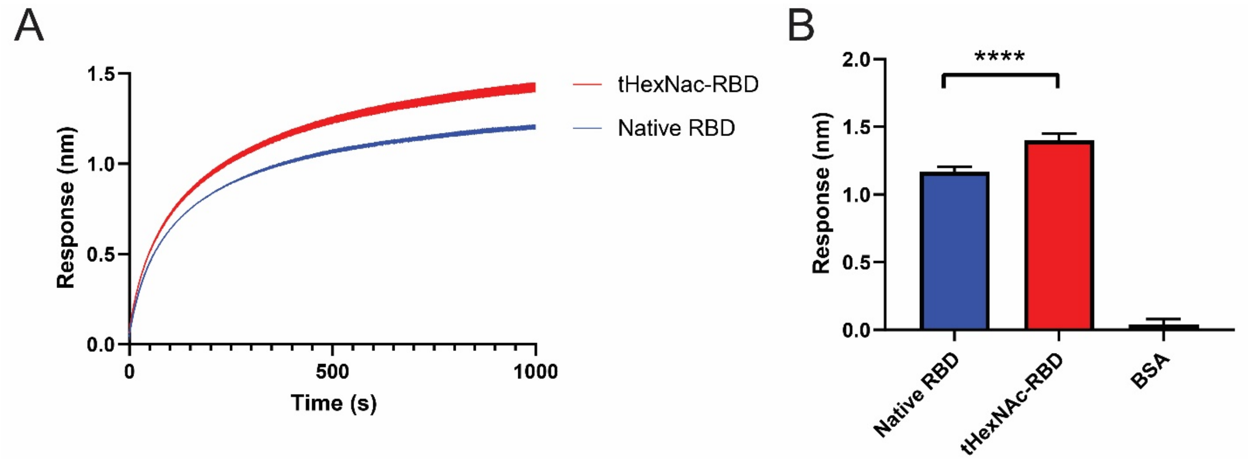


**Figure S4.** Comparison of ACE2 binding response (biolayer interferometry) between the native S protein RBD and tHexNAc RBD. (A) Response curve between 60 µg/ml ACE2 and 2 µg/ml RBD over 1000 seconds (Mean and SEM). (2) Comparison of ACE2 binding response at the 1000$^{th}$ second (Mean and SD). BSA was used as a negative control. Error bars: standard deviation (N=4); asterisks: P<0.0001.

Pr Proprietary

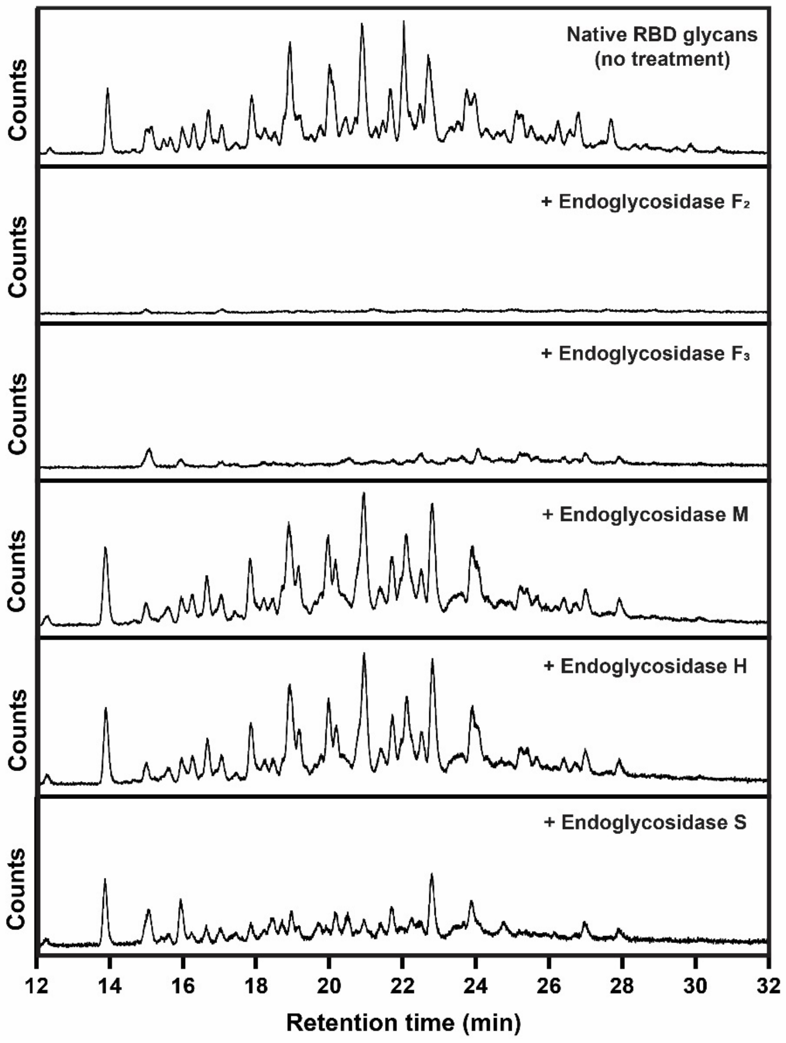


**Figure S5.** Activity screening of endoglycosidases on the SARs-CoV-2 S protein glycans. Active endoglycosidases are expected to remove N-glycans from the substrate, leading to a null signal in the analyses. Five glycosidase candidates were tested: Endoglycosidase F2 and F3 showed outstanding activity; while Endoglycosidase S had partial activity on the RBD glycans. Endoglycosidase M and H are known to be specific to high-mannose and hybrid-type N glycans that are in low content in the RBD glycan population. Reaction condition: 10% (w/w) enzyme-to-substrate for 24 hours at 37 °C. Endoglycosidases were purchased from NEB.

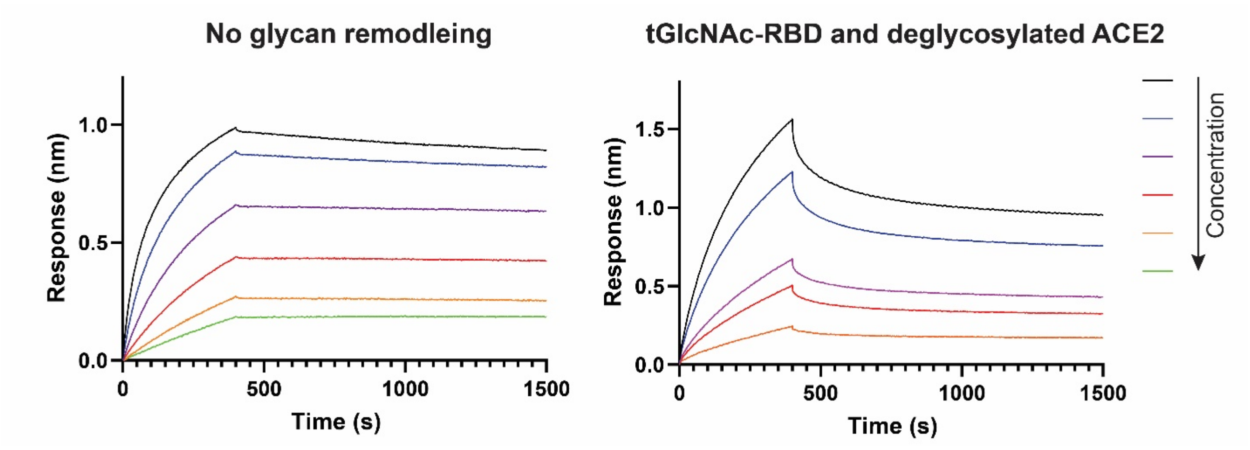


**Figure S6.** The removal of ACE2 glycans resulted in dramatic decrease of RBD binding affinity as presented by BLI response.

Pr Proprietary

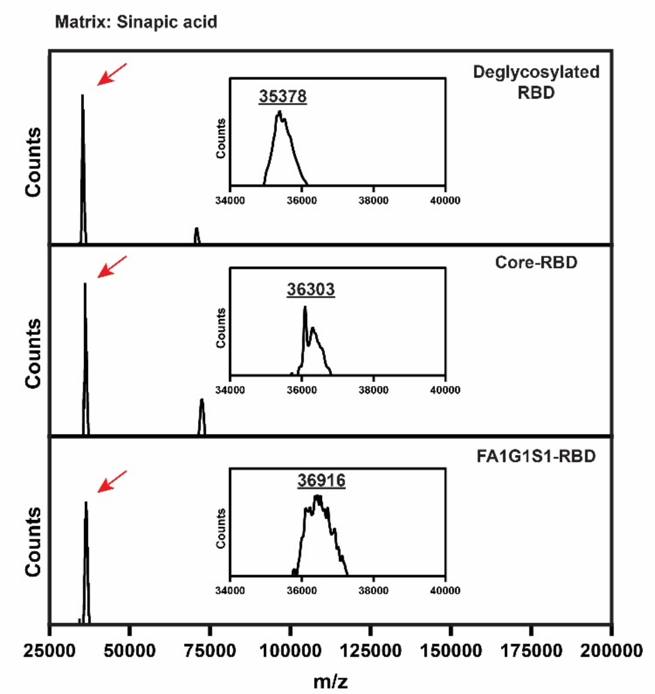


**Figure S7.** MALDI-ToF analysis supported the success of glycan structure remodeling on SARS-CoV-2 spike protein RBD.

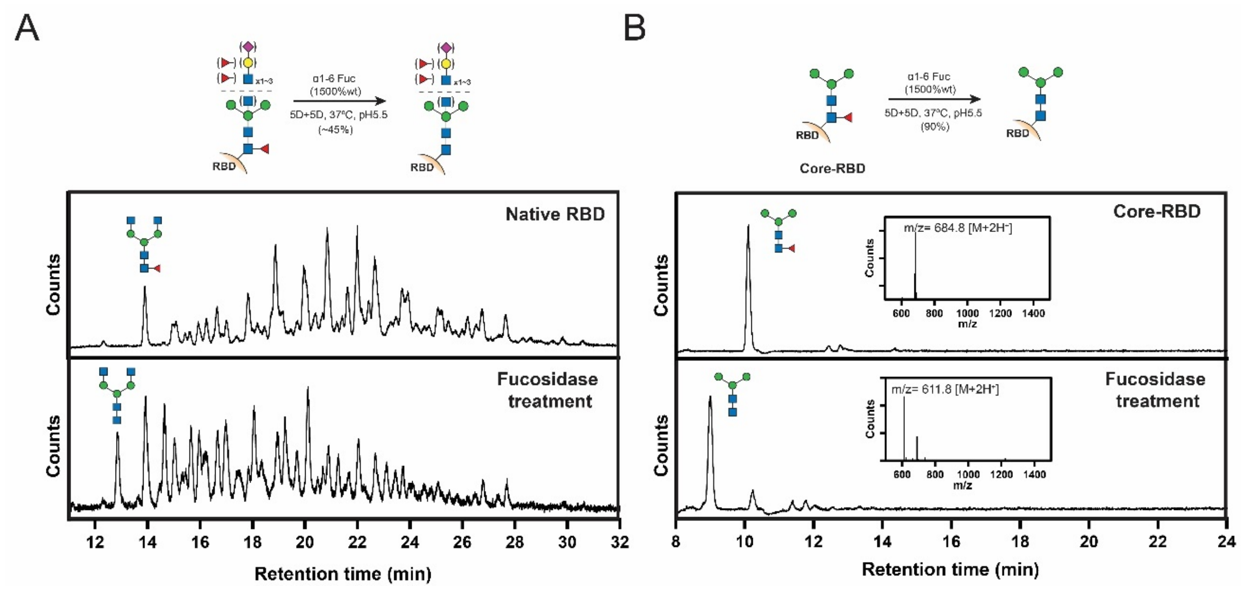


**Figure S8.** Removal of α1-6 fucose (core fucose) from SARS-CoV-2 S protein RBD. (A) α1-6 fucosidase treatment on native S protein RBD for 10 days. See SI-data. (B) α1-6 fucosidase treatment on the core-RBD for 10 days. Insertions are mass spectrum data collected from the highlighted peak. Note: isolated glycans were labeled with RapiFluor-MS before analyses.

Proprietary

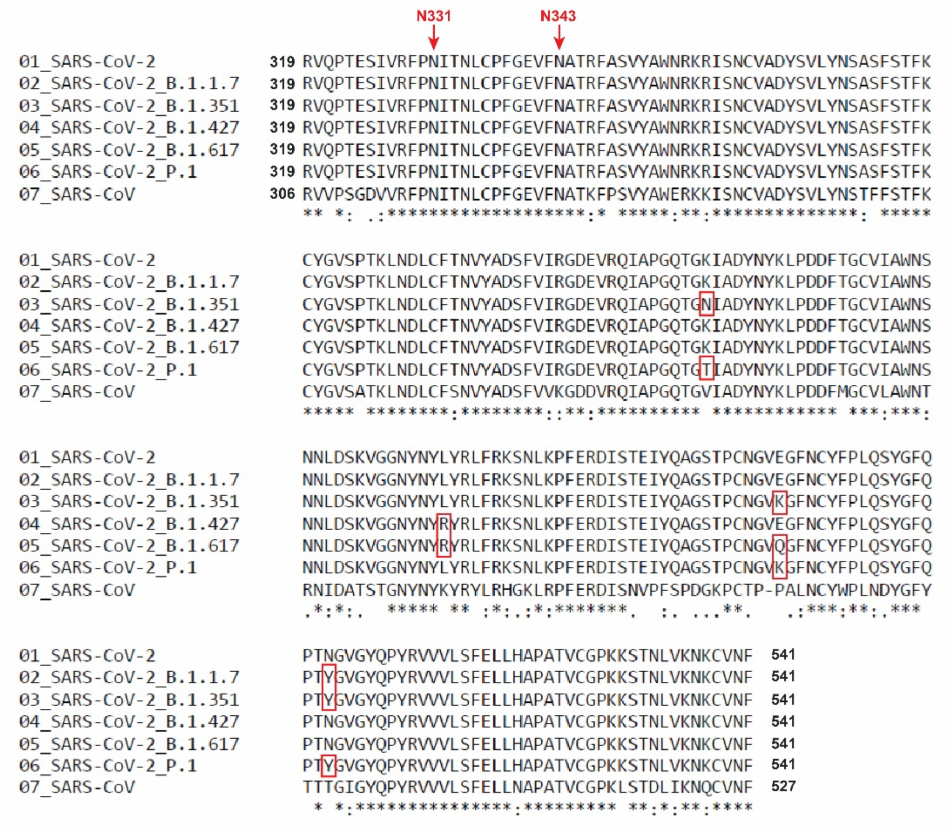


**Figure S9.** Sequence alignment of the receptor-binding domain (RBD) between SARS-CoV, SARS-CoV-2, and SARS-CoV-2 variants. The RBD glycosylation sites, N311 and N343, are highly conserved. Sequence information was published on the website of Centers for Disease Control and Prevention (CDC). Mutations found in the SARS-CoV-2 variants were highlighted by red boxes. The alignment was generated using Clustal Omega.(7)

Pr Proprietary

Proprietary
